# Robust computational design and evaluation of peptide vaccines for cellular immunity with application to SARS-CoV-2

**DOI:** 10.1101/2020.05.16.088989

**Authors:** Ge Liu, Brandon Carter, Trenton Bricken, Siddhartha Jain, Mathias Viard, Mary Carrington, David K. Gifford

## Abstract

We present a combinatorial machine learning method to evaluate and optimize peptide vaccine formulations, and we find for SARS-CoV-2 that it provides superior predicted display of viral epitopes by MHC class I and MHC class II molecules over populations when compared to other candidate vaccines. Our method is robust to idiosyncratic errors in the prediction of MHC peptide display and considers target population HLA haplotype frequencies during optimization. To minimize clinical development time our methods validate vaccines with multiple peptide presentation algorithms to increase the probability that a vaccine will be effective. We optimize an objective function that is based on the presentation likelihood of a diverse set of vaccine peptides conditioned on a target population HLA haplotype distribution and expected epitope drift. We produce separate peptide formulations for MHC class I loci (HLA-A, HLA-B, and HLA-C) and class II loci (HLA-DP, HLA-DQ, and HLA-DR) to permit signal sequence based cell compartment targeting using nucleic acid based vaccine platforms. Our SARS-CoV-2 MHC class I vaccine formulations provide 93.21% predicted population coverage with at least five vaccine peptide-HLA hits on average in an individual (≥ 1 peptide 99.91%) with all vaccine peptides perfectly conserved across 4,690 geographically sampled SARS-CoV-2 genomes. Our MHC class II vaccine formulations provide 90.17% predicted coverage with at least five vaccine peptide-HLA hits on average in an individual with all peptides having observed mutation probability ≤ 0.001. We evaluate 29 previously published peptide vaccine designs with our evaluation tool with the requirement of having at least five vaccine peptide-HLA hits per individual, and they have a predicted maximum of 58.51% MHC class I coverage and 71.65% MHC class II coverage given haplotype based analysis. We provide an open source implementation of our design methods (OptiVax), vaccine evaluation tool (EvalVax), as well as the data used in our design efforts.

## 1 Introduction

Peptide vaccines elicit a protective adaptive immune response to either cancer or infectious agent antigens to immunize against and combat ongoing disease [1, 2]. Their component peptides present undesired *epitopes* as 3D structural protein subunits or MHC displayed peptides to train the adaptive immune system to mount a response to a threat. T and B cells use their respective receptors to recognize vaccine presented epitopes to trigger activation and expansion of their response to the displayed epitopes. The activated and expanded T and B cells can then effectively mount a response against pathogens or tumor cells. Peptide vaccines are presently in development for cancer [3] and viral diseases including HIV [4], HCV, and Malaria [2, 5]. An HPV peptide vaccine is currently licensed for humans and encodes the sequence of two viral peptides that induce both CD4+ and CD8+ T cell responses [6].

The precise control of antigenic T cell recognized epitopes afforded by peptide vaccines has been proposed to reduce the risks posed by conventional vaccine approaches. For example, the conventional vaccine tetravalent dengue vaccine (CYD-TDV) increases the risk of hospitalization when an individual is infected with dengue for the first time. A study considered patients from 2 to 16 years of age that had not been infected at the time of vaccination but were infected post vaccination. The increased risk of hospitalization was thought to occur by antibody-dependent enhancement (ADE) by sub-neutralizing responses to the infecting dengue serotype [7]. A peptide based dengue vaccine has been proposed to induce CD4+ and CD8+ T cell response to dengue that would avoid ADE [8]. Given the multiple strains of coronavirus in circulation, considerations of ADE, immune enhancement, and other deleterious effects of vaccination need to be considered [9].

Here we focus on eliciting immunity by the adaptive immune system that is mediated by cells (cellular immunity). Cellular immunity can be induced with peptide vaccines that cause Major Histocompatibility Complex (MHC) molecules to display undesired epitopes on cell surfaces. Class I MHC molecules typically display peptides from a cell’s internal workings, while class II MHC molecules display peptides from a cell’s external environment that are taken up by professional antigen presenting cells by phagocytosis, and then made available for loading onto MHC class II molecules for cell surface display for T cell surveillance. CD8+ T cells recognize cells that are displaying non-self peptides on their class I MHC molecules and target the cells for destruction, while CD4+ T cells recognize non-self peptides on class II MHC molecules on professional antigen presenting cells and help prime the activation of CD8+ cells and antibody producing B cells. The production of a strong cellular immunity response to either a tumor or viral infection is important for positive patient outcomes. Cellular immunity is durable, and thus an important component of lasting immunity to viral infection.

There are multiple delivery platforms for peptide vaccines, including the direct injection of peptides in carriers and the delivery of recombinant nucleic acid that is turned into peptides by a patient’s cells. Recombinant nucleic acid delivery of vaccine formulations as either DNA or RNA has the advantage that it harnesses a patient’s own cells to transiently manufacture vaccine peptides. Recombinant nucleic acid vectors can be quickly adapted to new payloads. DNA or RNA can be delivered to cells via nanoparticles, non-pathogenic viruses, or other methods. DNA vaccines have the disadvantage that their DNA must be transported to the nucleus for transcription in mRNA. RNA vaccines can be delivered encapsulated in lipid nanoparticles that cells endocytose into the cytosol and translate into peptides [8, 10]. Peptides in a vaccine can be prepended with a signal sequence to stay within a cell’s cytosol for class I display, or be prepended with a different sequence to be transported the outside of a cell for class II display [11, 12]. A single mRNA molecule can be used to express class I and class II peptides with each class represented by an array of peptides separated by a 2A self-cleaving peptide site [13]. If desired, class II peptides can be fused to a protein subunit that is designed to elicit B cell responses and expressed in the same single mRNA molecule. In addition, class II peptides can be linked to Ii-Key peptides to enhance their presentation [14].

A challenge for the design of peptide vaccines is the diversity of human MHC alleles that each have specific preferences for the peptide sequences they will display. The Human Leukocyte Antigen (HLA) locus encodes the class I and class II MHC genes. We consider three loci that encode for MHC class I molecules (HLA-A, HLA-B, and HLA-C) and three loci that encode MHC class II molecules (HLA-DR, HLA-DQ, and HLA-DP). An individual’s HLA type describes the MHC alleles they contain at each of these loci. Peptides of length 8-10 residues can bind to MHC class I molecules whereas those of length 13-25 bind to MHC class II molecules [15, 16].

To create effective vaccines it is necessary to consider the MHC allelic frequency in the target population, as well as linkage disequilibrium between MHC genes to discover a set of peptides that is likely to be robustly displayed. Human populations that originate from different geographies have differing frequencies of MHC alleles, and these populations exhibit linkage disequilibrium between HLA loci that result in population specific haplotype frequencies. We utilize haplotype frequencies of three populations in the design and evaluation of our vaccine candidates.

Recent advances in machine learning have produced models that can predict the presentation of peptides by hundreds of allelic variants of both class I and class II MHC molecules [17, 18, 19, 20, 21]. These models are evaluated on their ability to accurately predict data unobserved during their training on hundreds of MHC alleles. Each method has its strengths and weaknesses. Given that different models may be more or less accurate for different sequence families and can make idiosyncratic errors, we use an ensemble of models for vaccine design. We evaluate completed designs using eleven models to provide a conservative evaluation of vaccine peptide presentation.

Previous peptide vaccine design and evaluation methods do not utilize the distribution of MHC haplotypes in a population, and thus can not accurately assess the coverage provided by a vaccine. These methods include VaxRank [22] that considers vaccine design for a single individual, and methods that do not take into account rare MHC allelic combinations including iVax [23], and SARS-CoV-2 specific efforts [24]. The IEDB Population Coverage Tool [25] estimates peptide-MHC binding coverage and the distribution of peptides displayed for a given population but assumes independence between different loci and thus does not consider linkage disequilibrium.

We consider methods for vaccine design within the following framework and assumptions. A method takes as input: the target proteome, the target proteome’s expected or observed conservation at amino acid resolution, and the target human population for vaccination, expressed in terms of the frequencies of their HLA haplotypes. A method outputs: a candidate set of MHC class I and a set of class II vaccine peptides. Target proteomes can be viral or oncogenes. Our methods eliminate peptides that are expected to be glycosylated, peptides that are expected to drift in sequence and thus cause vaccine escape, and peptides that are identical to peptides in the human proteome. Vaccine peptides can be drawn from the entire proteome or from specific proteins of interest. An overview of our system is shown in Figure 1.

**Figure 1.**
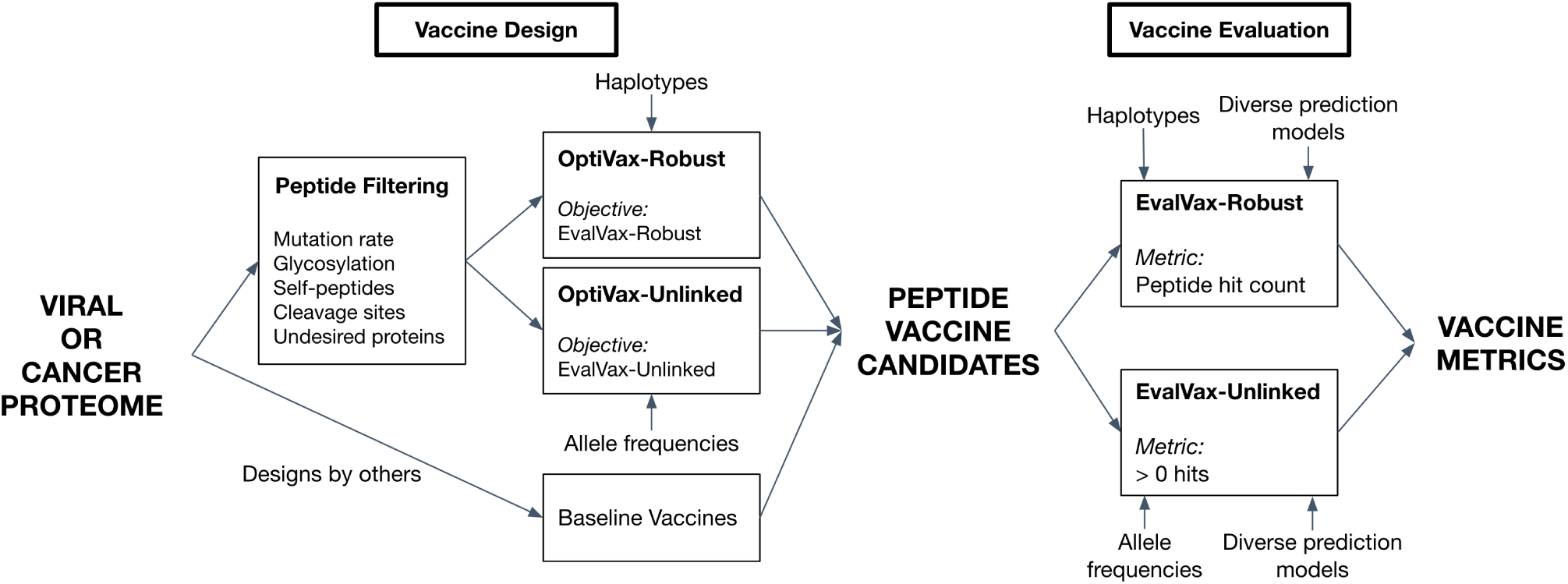
The OptiVax and EvalVax machine learning system for combinatorial vaccine optimization and evaluation.

We provide two methods for peptide vaccine evaluation, one that does not consider haplotype frequencies, EvalVax-Unlinked, and one that considers haplotype frequencies and computes the number of peptides predicted to be associated with population haplotypes, EvalVax-Robust. We employ these methods as objective functions for peptide vaccine formulation by combinatorial optimization in OptiVax-Unlinked and OptiVax-Robust. Using conservative metrics of peptide-MHC binding we find that our optimization methods provide both a higher likelihood of peptide display as well as a larger number of associated peptides than other published SARS-CoV-2 peptide vaccine designs with less than 150 peptides.

## 2 Methods

### 2.1 Datasets

#### A proteome is converted into candidate vaccine peptides

Given a target proteome as input, we identify all potential T cell epitopes for inclusion in a vaccine. We extract peptides of length 8-10 inclusive for consideration of MHC class I [15] binding and peptides of length 13-25 inclusive for class II [16] binding by using sliding windows of each size over the entire proteome. While peptides presented by MHC class I molecules can occasionally be longer than 10 residues [26], we conservatively limit our search to length 8-10 since MHC class I presented peptides are predominately 8-10 residues in length [15].

Using this sliding window approach, we created peptide sets from the SARS-CoV-2 (COVID-19) and SARS-CoV (Human SARS coronavirus) proteomes. SARS-CoV-2 was processed to discover relevant peptides for a vaccine, and SARS-CoV was processed to reveal common peptides between the two viruses during evaluation. The SARS-CoV-2 proteome is comprised of four structural proteins (E, M, N, and S) and at least six additional ORFs encoding nonstructural proteins, including the SARS-CoV-2 protease [27, 28]. We obtained the SARS-CoV-2 viral proteome from the GISAID [29] sequence entry Wuhan/IPBCAMS-WH-01/2019, the first documented case. We used Nextstrain [30] to identify open reading frames (ORFs) and translate the sequence. Our sliding windows on SARS-CoV-2 resulted in 29,403 candidate peptides for MHC class I and 125,593 candidate peptides for MHC class II. We obtained the SARS-CoV proteome from UniProt [31] under Proteome ID UP000000354. For SARS-CoV, our procedure creates 29,661 and 126,711 unique peptides for MHC class I and class II, respectively.

#### MHC population frequency computation

When we compute the probability of vaccine coverage over a population we use complementary methods that assume either independence or linkage between allele frequencies in genomically proximal HLA loci. In EvalVax-Unlinked (Section 2.4.2) we assume independence and use MHC allelic frequencies for 2392 class I alleles and 280 class II alleles from the dbMHC database [32] obtained from the IEDB Population Coverage Tool [25]. In EvalVax-Robust (Section 2.4.1) we assume linkage and use observed haplotype frequencies of HLA-A, HLA-B, and HLA-C loci for class I computations, or observed haplotype frequencies of HLA-DP, HLA-DQ, and HLA-DR for class II computations. We observed a total of 2138 distinct haplotypes for the HLA class I locus that include 230 different HLA-A, HLA-B, and HLA-C MHC alleles. We observed a total of 1711 distinct haplotypes for the HLA class II locus that include 280 different HLA-DP, HLA-DQ, and HLA-DR MHC alleles. We have independent haplotype frequency measurements for White, Black, and Asian populations.

HLA class I and class II haplotype frequencies were inferred using high resolution typing of individuals from distinct racial background. We estimated HLA class I haplotypes from HLA-A,-B, and -C genotypes of 2886 individuals of Black ancestry (46 distinct HLA-A alleles, 70 distinct HLA-B alleles, 40 distinct HLA-C alleles), 2327 individuals of White ancestry (38 distinct HLA-A alleles, 64 distinct HLA-B alleles, 34 distinct HLA-C alleles) and 1653 individuals of Asian ancestry (25 distinct HLA-A alleles, 51 distinct HLA-B alleles, 25 distinct HLA-C alleles). HLA class II haplotypes were estimated based on DR, DQ, DP genotypes of 2474 individuals of Black ancestry (10 distinct HLA-DPA1 alleles, 45 distinct HLA-DPB1 alleles, 14 distinct HLA-DQA1 alleles, 21 distinct HLA-DQB1 alleles, 38 distinct HLA-DRB1 alleles), 1857 individuals of White ancestry (7 distinct HLA-DPA1 alleles, 29 distinct HLA-DPB1 alleles, 18 distinct HLA-DQA1 alleles, 21 distinct HLA-DQB1 alleles, 41 distinct HLA-DRB1 alleles) and 1675 individuals of Asian ancestry (7 distinct HLA-DPA1 alleles, 28 distinct HLA-DPB1 alleles, 16 distinct HLA-DQA1 alleles, 16 distinct HLA-DQB1 alleles, 36 distinct HLA-DRB1 alleles). For each racial background, HLA class I and class II haplotypes were inferred using Hapferret [33] an implementation of the Expectation-Maximization algorithm [34]. A total of 1200, 779, and 440 class I and 920, 537, and 502 class II haplotype frequencies were derived in Black, White, and Asian populations, respectively.

### 2.2 Robust peptide-MHC binding prediction

#### Computational models

For a peptide vaccine to be effective, its constituent peptides need to be displayed, and thus a computational vaccine design must be built upon a solid predictive foundation of what peptides will be displayed by each MHC allele. Incorrect predictions could lead to failure of a pre-clinical or clinical trial at great human cost. To this end we are concerned with the precision (true positives / all positives) of our predictions such that we maximize the chance that a peptide predicted to be displayed will in fact be displayed. We are less concerned with our ability to recall all of the peptides that will work as long as we have a set of suitable size that will work. We reduce the risk of false positives by employing multiple computational methods to predict peptide-MHC binding. For design we use an ensemble of methods, and for evaluation we use all methods separately.

For MHC class I design, we use an ensemble that outputs the mean predicted binding affinity of NetMHCpan-4.0 [18] and MHCflurry 1.6.0 [35, 19]. We find this ensemble increases the precision of binding affinity estimates over the individual models on available SARS-CoV-2 experimental data (Table S1). For MHC class II design, we use NetMHCIIpan-4.0 [36]. For evaluation, we use our ensemble estimate of binding (MHC class I), as well as use binding predictions from a wide range of prediction algorithms (MHC class I: NetMHCpan-4.0 [18], NetMHCpan-4.1 [37], MHCflurry 1.6.0 [35], PUFFIN [17]; MHC class II: NetMHCIIpan-3.2 [20], NetMHCIIpan-4.0 [36], PUFFIN [17]) to ensure that all methods agree that we have a good peptide vaccine. We validate these models on datasets containing experimentally-studied SARS-CoV-2 and SARS-CoV peptides [38, 39, 40, 41] (see Section S1.2).

All models take as input a (MHC, peptide) pair and output predicted peptide-MHC binding affinity (IC50) on a nanomolar scale. For both MHC class I and class II models, we consider peptides to be binders if the predicted MHC binding affinity is ≤ 50nM [42]. This provides a conservative threshold to increase the probability of peptide display. Where our methods require a probability of peptide-MHC binding (as in Equation 5), affinity predictions are capped at 50000nM and transformed into [0, 1] using a logistic transformation, 1 − log_50000_(aff), where larger values correspond to greater likelihood of eliciting an immunogenic response [42, 43, 44]. The ≤ 50nM binding affinity threshold corresponds to a threshold of ≥ 0.638 after logistic transformation. We explored other criteria to classify peptides as binders and found using predicted binding affinity with a 50nM threshold to meet these alternative criteria and maximize precision on available SARS-CoV-2 experimental data (Table S1).

### 2.3 Removal of unfavorable peptides

#### 2.3.1 Removal of highly mutable peptides

We eliminate peptides that are observed to mutate above an input threshold rate to improve coverage over all SARS-CoV-2 variants and reduce the chance that the virus will mutate and escape vaccine-induced immunity in the future. When possible, we select peptides that are observed to be perfectly conserved across all observed SARS-CoV-2 viral genomes. Peptides that are observed to be perfectly conserved in thousands of examples may be functionally constrained to evolve slowly or not at all. If functional data are available, they can be used to supplement observed viral genome mutation rates by increasing mutation rates over functionally non-constrained residues.

For SARS-CoV-2, we obtained the most up to date version of the GISAID database [29] (as of 2:02pm EST May 13, 2020, acknowledgements in Section S4) and used Nextstrain [30] (from GitHub commit 639c63f25e0bf30c900f8d3d937de4063d96f791) to remove genomes with sequencing errors, translate the genome into proteins, and perform multiple sequence alignments (MSAs). We retrieved 24468 sequences from GISAID, and 19288 remained after Nextstrain quality processing. After quality processing, Nextstrain randomly sampled 34 genomes from every geographic region and month to produce a representative set of 5142 genomes for evolutionary analysis. Nextstrain definition of a “region” can vary from a city (e.g., “Shanghai”) to a larger geographical district. Spatial and temporal sampling in Nextstrain is designed to provide a representative sampling of sequences around the world.

The 5142 genomes sampled by Nextstrain were then translated into protein sequences and aligned. We eliminated viral genome sequences that had a stop codon, a gap, an unknown amino acid (because of an uncalled nucleotide in the codon), or had a gene that lacked a starting methionine, except for ORF1b which does not begin with a methionine. This left a total of 4690 sequences that were used to compute peptide level mutation probabilities. For each peptide, the probability of mutation was computed as the number of non-reference peptide sequences observed divided by the total number of peptide sequences observed.

#### 2.3.2 Removal of cleavage regions

SARS-CoV-2 contains a number of post-translation cleavage sites in ORF1a and ORF1b that result in a number of nonstructural protein products. Cleavage sites were obtained from UniProt [31] under entry P0DTD1. In addition, a furin-like cleavage site has been identified in the Spike protein [45]. This cleavage occurs before peptides are loaded in the endoplasmic reticulum for class I or endosomes for class II. Any peptide that spans any of these cleavage sites is removed from consideration. This removes 3,739 peptides out of the 154,996 we consider across windows 8-10 (class I) and 13-25 (class II) (∼2.4%).

#### 2.3.3 Removal of glycosylated peptides

We eliminate all peptides that are predicted to have N-linked glycosylation as it inhibits both MHC loading and T cell recognition of peptides [46]. Glycosylation is a post-translational modification that involves the covalent attachment of carbohydrates to specific motifs on the surface of the protein. We identified peptides that may be glycosylated with the NetNGlyc N-glycosylation prediction server [47]. We verified these predictions for the Spike protein using experimental data of Spike N-glycosylation from Cryo-EM and tandem mass spectrometry [48, 49]. A majority of the potential N-glycosylation sites (16 out of 22) were identified in both experimental studies, and further supported by homologous regions with glycosylation in SARS-CoV [50]. We found that that for the Spike protein when NetNGlyc predicted a non-zero probability of a site being N-glycosylated it was experimentally identified as a real or likely N-glycosylation site. Therefore, we eliminated all peptides where NetNGlyc predicted a non-zero N-glycosylation probability in any residue. This resulted in the elimination of 18,957 of the 154,996 peptides considered (∼12%).

#### 2.3.4 Self-epitope removal

T cells are selected to ignore peptides derived from the normal human proteome, and thus we remove any self peptides from consideration for a vaccine. In addition, it is possible that a vaccine might stimulate the adaptive immune system to react to a self peptide that was presented at an abnormally high level, which could lead to an autoimmune disorder. All peptides from SARS-CoV-2 were scanned against the entire human proteome downloaded from UniProt [31] under Proteome ID UP000005640. A total of 48 exact peptide matches (46 8-mers, two 9-mers) were discovered and eliminated from consideration.

#### 2.3.5 Removal of undesired proteins

OptiVax will design vaccines using peptides from specific viral or oncogene proteins of interest by removing peptides from undesired proteins from the candidate pool. Grifoni et al. [51] tested T cell responses from COVID-19 convalescent patients and found that peptides from the S, M, and N proteins of SARS-CoV-2 produce the dominant CD4+ and CD8+ responses when compared to other SARS-CoV-2 proteins. We have used OptiVax to produce additional SARS-CoV-2 vaccines comprised of peptides drawn from only S, M, and N as described in Section 3.2.

### 2.4 EvalVax evaluates peptide vaccine population coverage

We introduce two evaluation methods for estimating the population coverage of a proposed peptide vaccine set. EvalVax-Robust utilizes HLA haplotype frequencies for MHC class I (HLA-A/B/C) and MHC class II (HLA-DP/DQ/DR) genes, and evaluates population level likelihood of having larger than a certain number of peptide-HLA binding hits in each individual. EvalVax-Unlinked considers MHC allele frequencies at each HLA locus independently, and computes the likelihood that at least one peptide from a vaccine set is displayed at any locus. Both methods take into consideration MHC allele frequency, allelic zygosity, and for EvalVax-Robust, linkage disequilibrium (LD) among loci. We also take glycosylation and cleavage sites into consideration when evaluating vaccines by setting binding affinity to zero for peptides with non-zero glycosylation probability or on cleavage sites.

#### 2.4.1 EvalVax-Robust considers linkage disequilibrium of MHC genes

EvalVax-Robust computes the distribution of per individual peptide-HLA binding hits over a given population. It accounts for the significant linkage disequilibrium (LD) between HLA loci and uses haplotype frequencies for population coverage estimates. We expect that a vaccine will be more effective if more of its peptides are displayed by an individual’s MHC molecules, and thus EvalVax-Robust computes the probability of having at least *N* predicted peptide-HLA binding hits for each individual in the population.

Assuming for each of the HLA-A,B,C loci there are *M*_*A*_, *M*_*B*_, *M*_*C*_ alleles respectively, for a given haploid *A*_*i*_*B*_*j*_*C*_*k*_, the haplotype frequency is defined as *G*(*i, j, k*) and 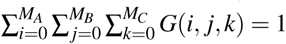. We assume independence of inherited haplotypes and compute the frequency of a diploid genotype as:

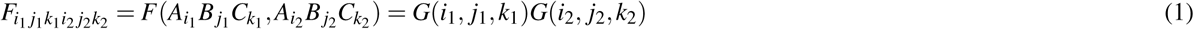

For each allele *A, e*(*A*) denotes the number of peptides predicted to bind to the allele with ≤ 50nM affinity, which we call the number of peptide-HLA hits. Then for each possible diploid genotype we compute the total number of peptide-HLA hits of the genotype as the sum of *e*(*A*) of the unique alleles in the genotype (there can be 3-6 unique alleles depending on the zygosity of each locus):

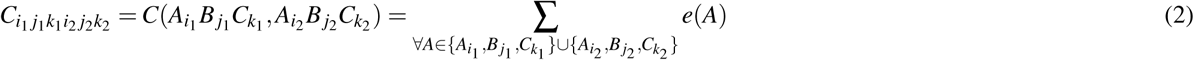

We then compute the frequency of having exactly *k* peptide-HLA hits in the population as:

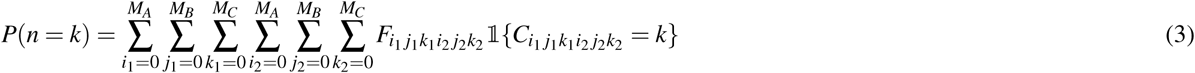

We define the population coverage objective function for EvalVax-Robust as the probability of having at least *N* peptide-HLA hits in the population, where the cutoff *N* is set to the minimum number of displayed vaccine peptides desired:

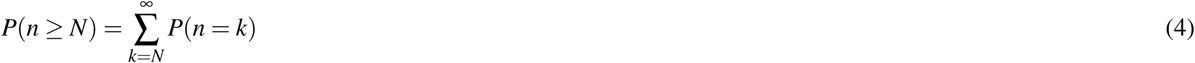

When we evaluate metrics on a world population, we equally weight population coverage estimations over three population groups (White, Black, and Asian) as the final objective function. In addition to the probability of having at least *N* peptide-HLA hits per individual, we also evaluate the expected number of per individual peptide-HLA hits in the population, which provides insight on how well the vaccine is displayed on average.

#### 2.4.2 EvalVax-Unlinked computes population coverage by at least one peptide-HLA hit

When haplotype frequencies are not available for a population, we can evaluate a vaccine using MHC allele frequencies that assume independence and compute the probability that at least one peptide binds to any of the alleles at any of the loci. To encourage a diverse set of peptides to bind to a single MHC allele, we use the predicted binding probability of a peptide to an allele instead of using a binary indicator of binding. This permits multiple peptides to contribute to the probability score at each allele. Considering *K* loci {*L*_1_, …, *L*_*K*_ }, for each locus there are *M*_*k*_ alleles 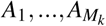 and the allele frequency is defined as *G*_*k*_(*A*_*i*_) and 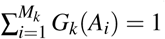. Given a set of *N* peptides {*P*_*n*=1:*N*_ }, for each allele (of locus *L*_*k*_) the predicted binding probability to peptide *P*_*n*_ is 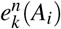. Assuming no competition between peptides, the probability that allele *A*_*i*_ ends up having at least one peptide bound is:

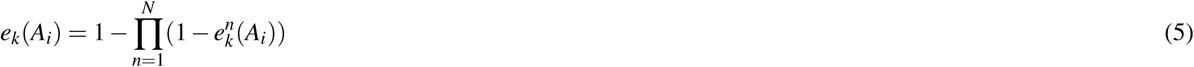

We define the diploid frequency of alleles as *F*_*k*_(*A*_*i*_, *A*_*j*_) = *G*_*k*_(*A*_*i*_)*G*_*k*_(*A*_*j*_), and we conservatively assume that a homozygous diploid locus does not improve the chance of peptide presentation over a single copy of the locus. Thus, the probability that a diploid genotype has at least one peptide bound is defined as:

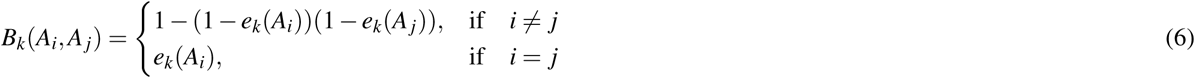

Therefore, the probability that a person in the given population displays at least one peptide in the set {*P*_*n*_ } at a particular locus *L*_*k*_ is calculated by:

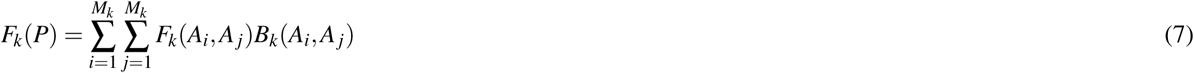

To combine different loci assuming no linkage disequilibrium, the probability that a person in the given population has at least one locus that binds to at least one peptide from {*P*_*n*_} is defined as:

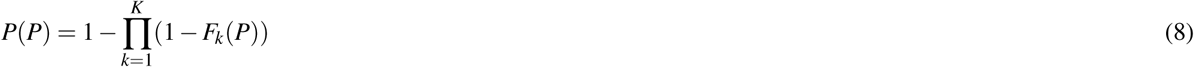

which is the evaluation metric for EvalVax-Unlinked.

We conservatively only consider peptides with predicted binding affinity ≤ 50nM. We set values of 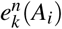 weaker than 50nM predicted binding affinity to zero. This constraint on peptide binding is in addition to all of the other peptide filters in Section 2.3. When we evaluate on a world population, we equally weight population coverage estimates over 15 geographic regions (see Results for list of regions) as the final objective function.

### 2.5 OptiVax selects optimized vaccine peptide sets

We use beam search over a set of candidate peptides to efficiently search for an optimal subset of peptides that maximizes a desired EvalVax-Unlinked or EvalVal-Robust based objective function. Our beam search procedure is parallelizable across CPU cores, and we typically use from 40 to 96 cores. We use a beam size of *k* = 10 for MHC class I and *k* = 5 for MHC class II.

#### 2.5.1 OptiVax-Robust searches for a peptide set with high expected number of per-individual peptide-HLA hits

OptiVax-Robust uses beam search to find a minimal set of peptides that reaches a desired population coverage probability at a threshold of *N* predicted peptide-HLA hits for each individual. We start from an empty set of peptides and *N* = 0, and iteratively expand the solution by one peptide at a time and retain the top *k* solutions until the population coverage probability for the current *N* reaches the given population coverage probability threshold for that *N*. We then repeat the same process for *N* + 1. At the expense of increased computational cost, beam search improves upon greedy optimization by considering *k* possible solutions at each step. During each iteration, the population coverage probability threshold at the present *N* controls the robustness of coverage. Increasing the desired population coverage probability increases the difficulty of the optimization task. The iterative process stops when a desired population coverage at a desired *N* is achieved. In early rounds of optimization, OptiVax uses a high population coverage probability to provide better individual coverage. In subsequent rounds, the target population coverage probability is reduced on a fixed schedule.

#### 2.5.2 OptiVax-Unlinked searches for a peptide set that covers a population

OptiVax-Unlinked uses beam search to find a minimal set of peptides that reaches a desired population coverage probability that each individual on average displays at least one vaccine peptide. We iteratively expand solutions in the beam by adding one peptide at a time to reach the population coverage objective, and keep the top *k* solutions over all possible expansions in the beam.

#### 2.5.3 OptiVax improves vaccine sequence diversity

OptiVax reduces vaccine sequence redundancy by not selecting peptides with closely related sequences for a vaccine formulation. This issue arises because sliding a window over a proteome produces overlapping sequences that are very similar in MHC binding characteristics. When any version of OptiVax selects a peptide during optimization, it eliminates from further consideration all unselected peptides that are within three (MHC class I) or five (MHC class II) edits on a sequence distance metric from the selected peptide. The distance metric aligns two peptides without gaps within them and is the sum of the lengths of their unaligned portions at their ends.

## 3 Results

### 3.1 Validation of peptide-MHC binding prediction models for OptiVax design

We validate our computational models on datasets containing experimentally-studied SARS-CoV-2 and SARS-CoV peptides [38, 39, 40, 41] (details in Section S1.2). We find classifying peptides as binders by predicted binding affinity ≤ 50nM maximizes AUROC and precision in classification of stable binders over alternative predictors and binding criteria (Table S1). Our ensemble of NetMHCpan-4.0 and MHCflurry further increases AUROC and precision over individual predictors.

### 3.2 OptiVax-Robust optimization results on MHC class I and II

#### MHC class I results

We selected an optimized set of peptides from all SARS-CoV-2 proteins using the EvalVax-Robust objective function. We limited our candidates to peptides with length 8-10 and excluded peptides that have been observed with any mutation or are predicted to have non-zero probability of glycosylation. For computation of the objective function, we use the mean predicted IC50 values from our NetMHCpan-4.0 and MHCflurry ensemble to obtain reliable binding affinity predictions for evaluation and optimization. With OptiVax-Robust optimization, we design a vaccine with 19 peptides that achieves 99.39% EvalVax-Unlinked coverage and 99.91% EvalVax-Robust coverage over three ethnic groups (Asian, Black, White) with at least one peptide-HLA hit per individual. This set of peptides also provides 93.21% coverage with at least 5 peptide-HLA hits and 67.75% coverage with at least 8 peptide-HLA hits (Figure 2, Table 1). The population level distribution of the number of peptide-HLA hits in White, Black, and Asian populations is shown in Figure 2, where the expected number of peptide-HLA hits is 9.358, 8.515, and 10.206, respectively.

**Table 1.**
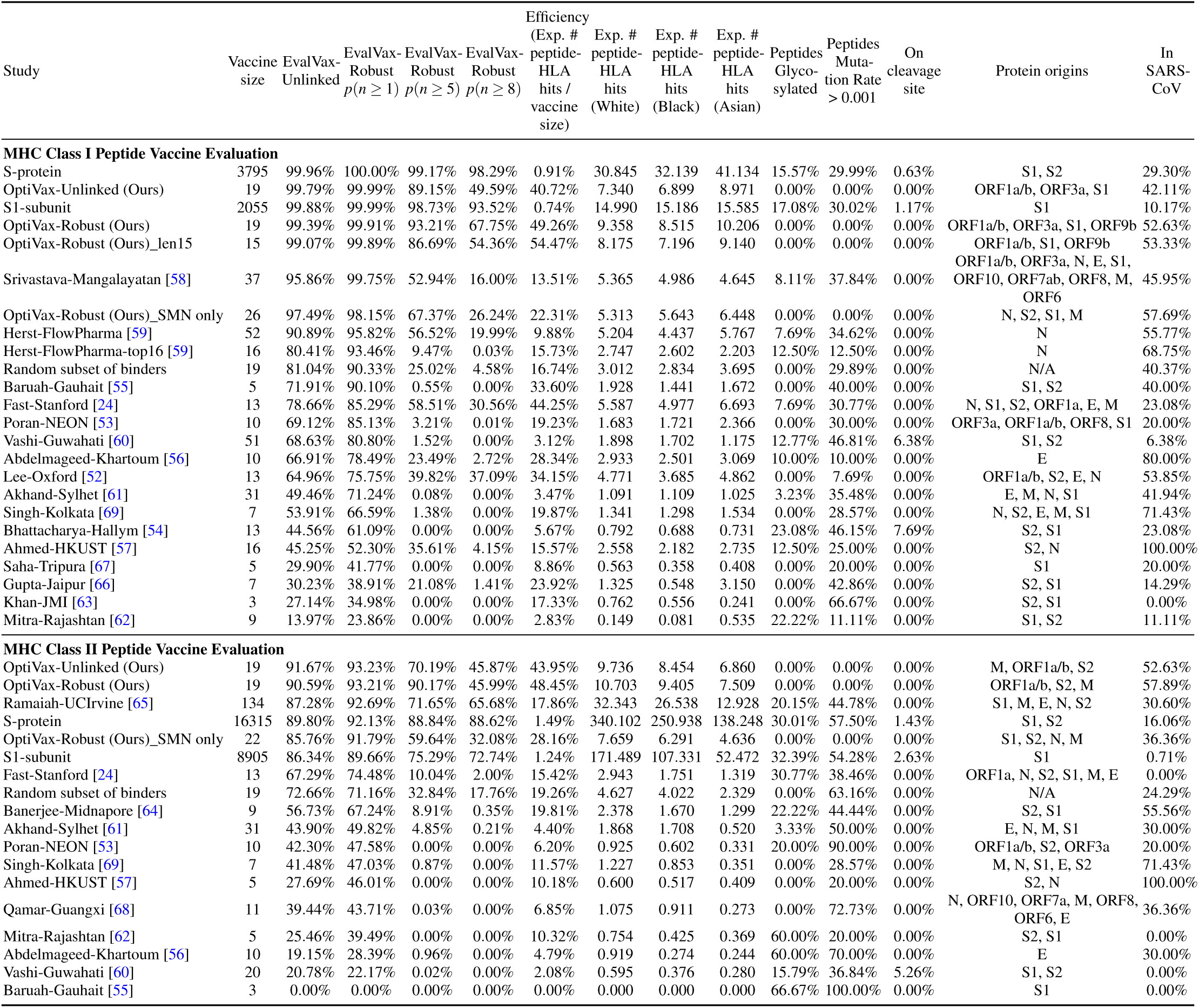
Comparison of existing baselines, S-protein peptides, and OptiVax designed peptide vaccines (using full set of proteins or S/M/N proteins only) on various population coverage evaluation metrics and vaccine quality metrics (percentage of peptides with larger than 0.1% probability of mutating or with non-zero probability of being glycosylated). The list is sorted by EvalVax-Robust *p*(*n* ≥ 1). Random subsets are generated 200 times.The binders used for generating random subsets are defined as peptides that is predicted to bind with ≤ 50nM to more than 5 of the alleles.

| Study | Vaccine size | EvalVax-Unlinked | EvalVax-Robust<br>$p(n \geq 1)$ | EvalVax-Robust<br>$p(n \geq 5)$ | EvalVax-Robust<br>$p(n \geq 8)$ | Efficiency<br>(Exp. #<br>peptide-HLA<br>hits /<br>vaccine<br>size) | Exp. #<br>peptide-HLA<br>hits<br>(White) | Exp. #<br>peptide-HLA<br>hits<br>(Black) | Exp. #<br>peptide-HLA<br>hits<br>(Asian) | Peptides<br>Glyco-<br>sylated | Peptides<br>Muta-<br>tion Rate<br>> 0.001 | On<br>cleavage<br>site | Protein origins | In<br>SARS-CoV |
| --- | --- | --- | --- | --- | --- | --- | --- | --- | --- | --- | --- | --- | --- | --- |
| **MHC Class I Peptide Vaccine Evaluation** |  |  |  |  |  |  |  |  |  |  |  |  |  |  |
| S-protein | 3795 | 99.96% | 100.00% | 99.17% | 98.29% | 0.91% | 30.845 | 32.139 | 41.134 | 15.57% | 29.99% | 0.63% | S1, S2 | 29.30% |
| OptiVax-Unlinked (Ours) | 19 | 99.79% | 99.99% | 89.15% | 49.59% | 40.72% | 7.340 | 6.899 | 8.971 | 0.00% | 0.00% | 0.00% | ORF1a/b, ORF3a, S1 | 42.11% |
| S1-subunit | 2055 | 99.88% | 99.99% | 98.73% | 93.52% | 0.74% | 14.990 | 15.186 | 15.585 | 17.08% | 30.02% | 1.17% | S1 | 10.17% |
| OptiVax-Robust (Ours) | 19 | 99.39% | 99.91% | 93.21% | 67.75% | 49.26% | 9.358 | 8.515 | 10.206 | 0.00% | 0.00% | 0.00% | ORF1a/b, ORF3a, S1, ORF9b | 52.63% |
| OptiVax-Robust (Ours)_len15 | 15 | 99.07% | 99.89% | 86.69% | 54.36% | 54.47% | 8.175 | 7.196 | 9.140 | 0.00% | 0.00% | 0.00% | ORF1a/b, S1, ORF9b | 53.33% |
| Srivastava-Mangalayatan [58] | 37 | 95.86% | 99.75% | 52.94% | 16.00% | 13.51% | 5.365 | 4.986 | 4.645 | 8.11% | 37.84% | 0.00% | ORF1a/b, ORF3a, N, E, S1,<br>ORF10, ORF7ab, ORF8, M,<br>ORF6 | 45.95% |
| OptiVax-Robust (Ours)_SMN only | 26 | 97.49% | 98.15% | 67.37% | 26.24% | 22.31% | 5.313 | 5.643 | 6.448 | 0.00% | 0.00% | 0.00% | N, S2, S1, M | 57.69% |
| Herst-FlowPharma [59] | 52 | 90.89% | 95.82% | 56.52% | 19.99% | 9.88% | 5.204 | 4.437 | 5.767 | 7.69% | 34.62% | 0.00% | N | 55.77% |
| Herst-FlowPharma-top16 [59] | 16 | 80.41% | 93.46% | 9.47% | 0.03% | 15.73% | 2.747 | 2.602 | 2.203 | 12.50% | 12.50% | 0.00% | N | 68.75% |
| Random subset of binders | 19 | 81.04% | 90.33% | 25.02% | 4.58% | 16.74% | 3.012 | 2.834 | 3.695 | 0.00% | 29.89% | 0.00% | N/A | 40.37% |
| Baruah-Gauhait [55] | 5 | 71.91% | 90.10% | 0.55% | 0.00% | 33.60% | 1.928 | 1.441 | 1.672 | 0.00% | 40.00% | 0.00% | S1, S2 | 40.00% |
| Fast-Stanford [24] | 13 | 78.66% | 85.29% | 58.51% | 30.56% | 44.25% | 5.587 | 4.977 | 6.693 | 7.69% | 30.77% | 0.00% | N, S1, S2, ORF1a, E, M | 23.08% |
| Poran-NEON [53] | 10 | 69.12% | 85.13% | 3.21% | 0.01% | 19.23% | 1.683 | 1.721 | 2.366 | 0.00% | 30.00% | 0.00% | ORF3a, ORF1a/b, ORF8, S1 | 20.00% |
| Vashi-Guwahati [60] | 51 | 68.63% | 80.80% | 1.52% | 0.00% | 3.12% | 1.898 | 1.702 | 1.175 | 12.77% | 46.81% | 6.38% | S1, S2 | 6.38% |
| Abdelmageed-Khartoum [56] | 10 | 66.91% | 78.49% | 23.49% | 2.72% | 28.34% | 2.933 | 2.501 | 3.069 | 10.00% | 10.00% | 0.00% | E | 80.00% |
| Lee-Oxford [52] | 13 | 64.96% | 75.75% | 39.82% | 37.09% | 34.15% | 4.771 | 3.685 | 4.862 | 0.00% | 7.69% | 0.00% | ORF1a/b, S2, E, N | 53.85% |
| Akhand-Sylhet [61] | 31 | 49.46% | 71.24% | 0.08% | 0.00% | 3.47% | 1.091 | 1.109 | 1.025 | 3.23% | 35.48% | 0.00% | E, M, N, S1 | 41.94% |
| Singh-Kolkata [69] | 7 | 53.91% | 66.59% | 1.38% | 0.00% | 19.87% | 1.341 | 1.298 | 1.534 | 0.00% | 28.57% | 0.00% | N, S2, E, M, S1 | 71.43% |
| Bhattacharya-Hallym [54] | 13 | 44.56% | 61.09% | 0.00% | 0.00% | 5.67% | 0.792 | 0.688 | 0.731 | 23.08% | 46.15% | 7.69% | S2, S1 | 23.08% |
| Ahmed-HKUST [57] | 16 | 45.25% | 52.30% | 35.61% | 4.15% | 15.57% | 2.558 | 2.182 | 2.735 | 12.50% | 25.00% | 0.00% | S2, N | 100.00% |
| Saha-Tripura [67] | 5 | 29.90% | 41.77% | 0.00% | 0.00% | 8.86% | 0.563 | 0.358 | 0.408 | 0.00% | 20.00% | 0.00% | S1 | 20.00% |
| Gupta-Jaipur [66] | 7 | 30.23% | 38.91% | 21.08% | 1.41% | 23.92% | 1.325 | 0.548 | 3.150 | 0.00% | 42.86% | 0.00% | S2, S1 | 14.29% |
| Khan-JMI [63] | 3 | 27.14% | 34.98% | 0.00% | 0.00% | 17.33% | 0.762 | 0.556 | 0.241 | 0.00% | 66.67% | 0.00% | S2, S1 | 0.00% |
| Mitra-Rajasthan [62] | 9 | 13.97% | 23.86% | 0.00% | 0.00% | 2.83% | 0.149 | 0.081 | 0.535 | 22.22% | 11.11% | 0.00% | S1, S2 | 11.11% |
| **MHC Class II Peptide Vaccine Evaluation** |  |  |  |  |  |  |  |  |  |  |  |  |  |  |
| OptiVax-Unlinked (Ours) | 19 | 91.67% | 93.23% | 70.19% | 45.87% | 43.95% | 9.736 | 8.454 | 6.860 | 0.00% | 0.00% | 0.00% | M, ORF1a/b, S2 | 52.63% |
| OptiVax-Robust (Ours) | 19 | 90.59% | 93.21% | 90.17% | 45.99% | 48.45% | 10.703 | 9.405 | 7.509 | 0.00% | 0.00% | 0.00% | ORF1a/b, S2, M | 57.89% |
| Ramaiah-UCIrvine [65] | 134 | 87.28% | 92.69% | 71.65% | 65.68% | 17.86% | 32.343 | 26.538 | 12.928 | 20.15% | 44.78% | 0.00% | S1, M, E, N, S2 | 30.60% |
| S-protein | 16315 | 89.80% | 92.13% | 88.84% | 88.62% | 1.49% | 340.102 | 250.938 | 138.248 | 30.01% | 57.50% | 1.43% | S1, S2 | 16.06% |
| OptiVax-Robust (Ours)_SMN only | 22 | 85.76% | 91.79% | 59.64% | 32.08% | 28.16% | 7.659 | 6.291 | 4.636 | 0.00% | 0.00% | 0.00% | S1, S2, N, M | 36.36% |
| S1-subunit | 8905 | 86.34% | 89.66% | 75.29% | 72.74% | 1.24% | 171.489 | 107.331 | 52.472 | 32.39% | 54.28% | 2.63% | S1 | 0.71% |
| Fast-Stanford [24] | 13 | 67.29% | 74.48% | 10.04% | 2.00% | 15.42% | 2.943 | 1.751 | 1.319 | 30.77% | 38.46% | 0.00% | ORF1a, N, S2, S1, M, E | 0.00% |
| Random subset of binders | 19 | 72.66% | 71.16% | 32.84% | 17.76% | 19.26% | 4.627 | 4.022 | 2.329 | 0.00% | 63.16% | 0.00% | N/A | 24.29% |
| Banerjee-Midnapore [64] | 9 | 56.73% | 67.24% | 8.91% | 0.35% | 19.81% | 2.378 | 1.670 | 1.299 | 22.22% | 44.44% | 0.00% | S2, S1 | 55.56% |
| Akhand-Sylhet [61] | 31 | 43.90% | 49.82% | 4.85% | 0.21% | 4.40% | 1.868 | 1.708 | 0.520 | 3.33% | 50.00% | 0.00% | E, N, M, S1 | 30.00% |
| Poran-NEON [53] | 10 | 42.30% | 47.58% | 0.00% | 0.00% | 6.20% | 0.925 | 0.602 | 0.331 | 20.00% | 90.00% | 0.00% | ORF1a/b, S2, ORF3a | 20.00% |
| Singh-Kolkata [69] | 7 | 41.48% | 47.03% | 0.87% | 0.00% | 11.57% | 1.227 | 0.853 | 0.351 | 0.00% | 28.57% | 0.00% | M, N, S1, E, S2 | 71.43% |
| Ahmed-HKUST [57] | 5 | 27.69% | 46.01% | 0.00% | 0.00% | 10.18% | 0.600 | 0.517 | 0.409 | 0.00% | 20.00% | 0.00% | S2, N | 100.00% |
| Qamar-Guangxi [68] | 11 | 39.44% | 43.71% | 0.03% | 0.00% | 6.85% | 1.075 | 0.911 | 0.273 | 0.00% | 72.73% | 0.00% | N, ORF10, ORF7a, M, ORF8,<br>ORF6, E | 36.36% |
| Mitra-Rajasthan [62] | 5 | 25.46% | 39.49% | 0.00% | 0.00% | 10.32% | 0.754 | 0.425 | 0.369 | 60.00% | 20.00% | 0.00% | S2, S1 | 0.00% |
| Abdelmageed-Khartoum [56] | 10 | 19.15% | 28.39% | 0.96% | 0.00% | 4.79% | 0.919 | 0.274 | 0.244 | 60.00% | 70.00% | 0.00% | E | 30.00% |
| Vashi-Guwahati [60] | 20 | 20.78% | 22.17% | 0.02% | 0.00% | 2.08% | 0.595 | 0.376 | 0.280 | 15.79% | 36.84% | 5.26% | S1, S2 | 0.00% |
| Baruah-Gauhait [55] | 3 | 0.00% | 0.00% | 0.00% | 0.00% | 0.00% | 0.000 | 0.000 | 0.000 | 66.67% | 100.00% | 0.00% | S1 | 0.00% |

**Figure 2.**
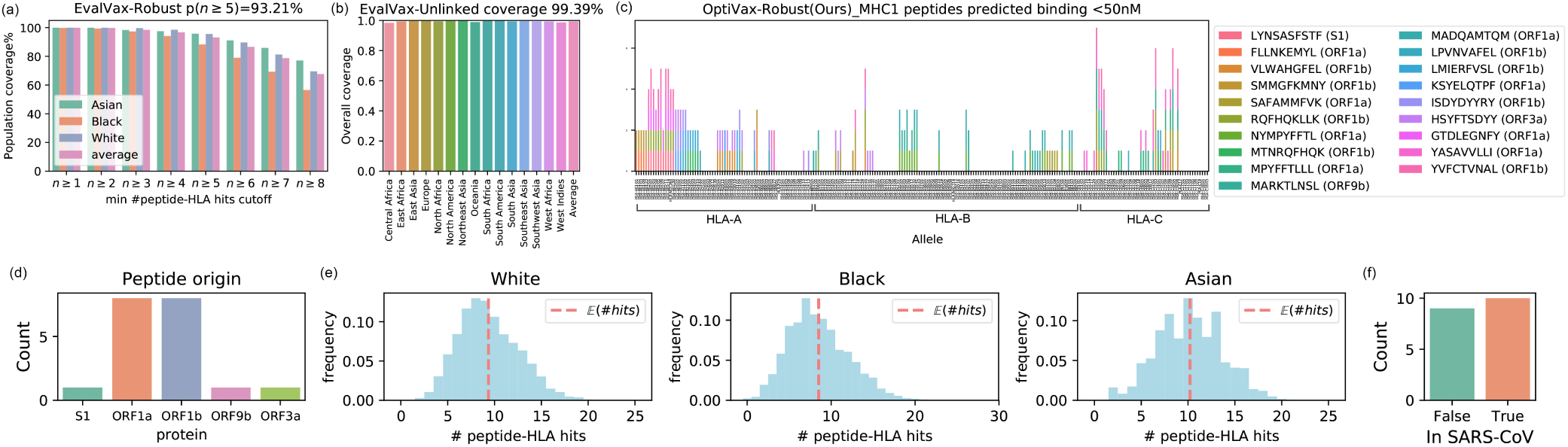
OptiVax-Robust selected peptide set for MHC class I. (a) EvalVax-Robust population coverage at different per-individual number of peptide-HLA hit cutoffs for Asian/Black/White populations and average value. (b) EvalVax-Unlinked population coverage on 15 geographic regions and averaged population coverage. (c) Binding of vaccine peptides to 230 HLA-A/B/C alleles. (d) Distribution of peptide origin. (e) Distribution of the number of per-individual peptide-HLA hits in White/Black/Asian populations. (f) Peptide presence in SARS-CoV.

#### MHC class II results

We limited our candidates to peptides with length 13-25 and excluded peptides that have been observed with mutation probability greater than 0.001 or are predicted to have non-zero glycosylation probability. We use the predicted binding affinity from NetMHCIIpan-4.0 for optimization and evaluation. With OptiVax-Robust optimization, we design a vaccine with 20 peptides that achieves 90.59% EvalVax-Unlinked coverage and 93.21% EvalVax-Robust coverage over three ethnic groups (Asian, Black, White) with at least one peptide-HLA hit per individual. This set of peptides also provides 90.17% coverage with at least 5 peptide-HLA hits and 45.99% coverage with at least 8 peptide-HLA hits (Figure 3, Table 1). The population level distribution of the number of peptide-HLA hits per individual in White, Black, and Asian populations is shown in Figure 3, where the expected number of of peptide-HLA hits is 10.703, 9.405, and 7.509, respectively.

**Figure 3.**
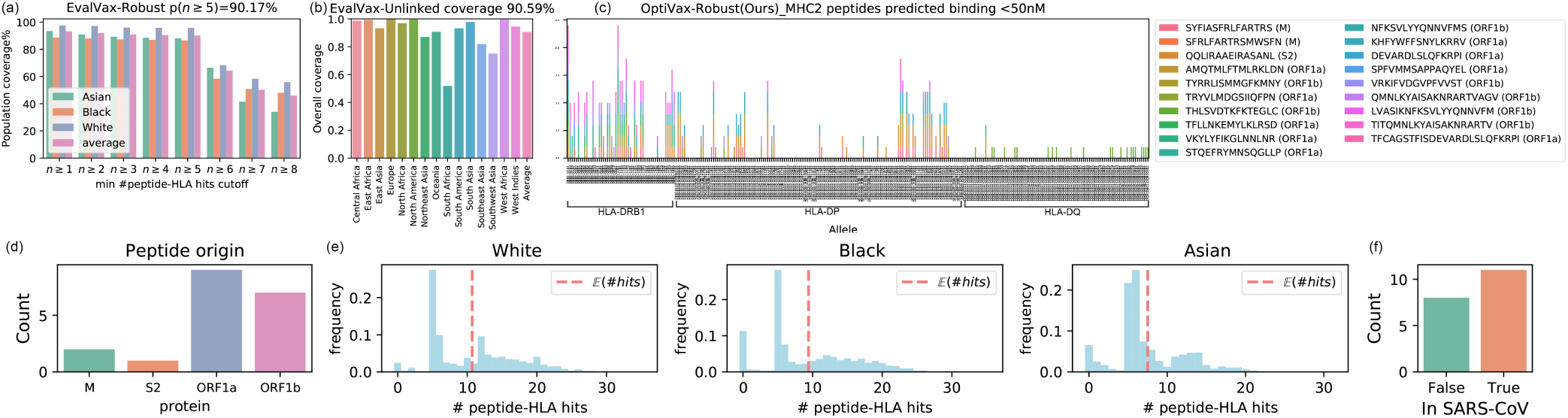
OptiVax-Robust selected optimal peptide set for MHC class II. (a) EvalVax-Robust population coverage at different minimum number of peptide-HLA hit cutoffs. (b) EvalVax-Unlinked population coverage. (c) Binding of vaccine peptides to 280 HLA-DRB1/DP/DQ alleles. (d) Distribution of peptide origin. (e) Distribution of the number of per-individual peptide-HLA hits in White/Black/Asian populations. (f) Peptide presence in SARS-CoV.

#### Designing vaccines with S, M, N proteins only

We also used OptiVax-Robust to design vaccines for MHC class I and class II based solely upon peptides from the S, M, and N proteins of SARS-CoV-2 and evaluated vaccine performance. Grifoni et al. [51] found that peptides from the S, M, and N structural proteins of SARS-CoV-2 were dominant in producing responses from CD4+ and CD8+ cells from convalescent COVID-19 patients. As shown in Table 1, the resulting MHC class I vaccine with 26 peptides achieves 98.15% coverage over three ethnic groups (Asian, Black, White) with at least one average peptide-HLA hit per individual. There were an average of at least five peptide hits in 67.37% of the population, and the expected per-individual number of hits for White, Black, and Asian populations are 5.313, 5.643, and 6.448, respectively. The OptiVax-Robust MHC class II vaccine with 22 S, M, and N peptides achieves 91.79% coverage with an average of at least one peptide-HLA hit per individual. There were an average of at least five peptide hits in 59.64% of the population, and the expected per-individual number of hits in White, Black, and Asian populations are 7.659, 6.291, and 4.636, respectively. The detailed vaccine designs are in Figure S1. We observed that it is more difficult to optimize vaccines with S, N, and M proteins only. We expect this is because we have fewer candidate peptides to cover all of our haplotype combinations.

### 3.3 OptiVax-Unlinked optimization results on MHC class I and II

#### MHC class I results

We limited our candidates to peptides with length 8-10 and zero predicted probability of glycosylation. We also excluded peptides that have been observed with any mutation. We use the mean predicted binding affinity values from our ensemble of NetMHCpan-4.0 and MHCflurry on 2392 MHC class I alleles to obtain reliable binding affinity predictions for evaluation and optimization. With OptiVax-Unlinked optimization, we design a vaccine with 19 peptides that achieves 99.79% EvalVax-Unlinked population coverage (averages over 15 geographic regions). As shown in Figure 4, the 19 vaccine peptides bind to a diverse range of alleles across the HLA-A/B/C loci. Even though less effective than OptiVax-Robust at providing a higher number of expected individual peptide-HLA hits in the population, the OptiVax-Unlinked peptide set still achieves high coverage on EvalVax-Robust metrics (99.99% for *p*(*n* ≥ 1), 89.15% for *p*(*n* ≥ 5), 49.59% for *p*(*n* ≥ 8)). The expected per-individual number of peptide-HLA hits for the design is 7.340, 6.899, and 8.971 for White, Black, and Asian populations, respectively (Table 1).

**Figure 4.**
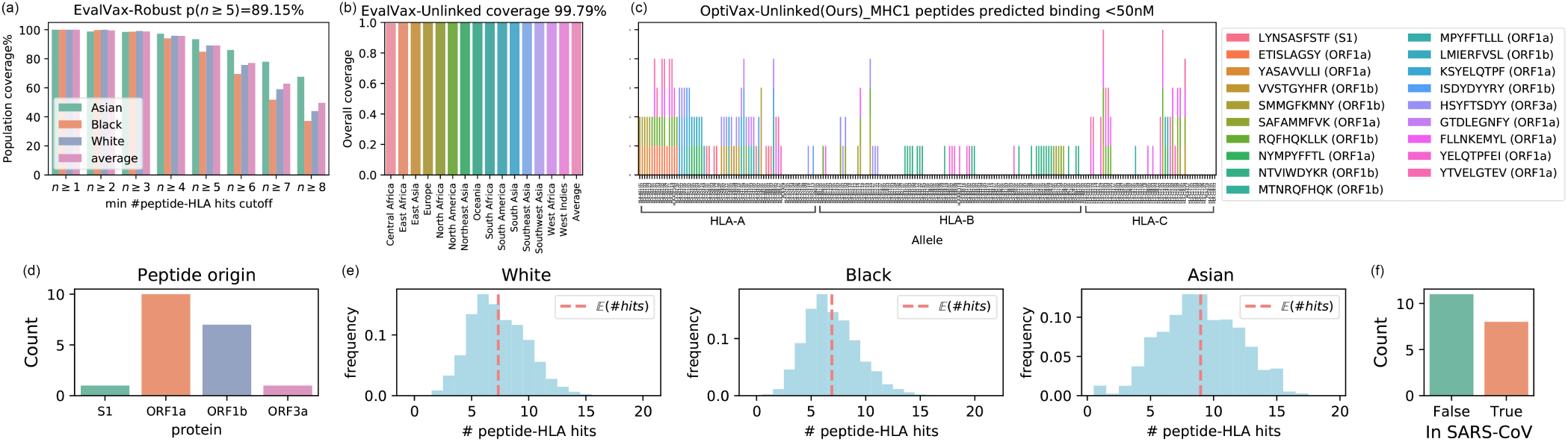
OptiVax-Unlinked selected optimal peptide set for MHC class I. (a) EvalVax-Robust population coverage at different per-individual number of peptide-HLA hits cutoffs for Asian/Black/White populations and average value. (b) EvalVax-Unlinked population coverage on 15 geographic regions and averaged population coverage. (c) Binding of vaccine peptides to 230 HLA-A/B/C alleles. (d) Distribution of peptide origin. (e) Distribution of the number of per-individual peptide-HLA hits in White/Black/Asian populations. (f) Peptide presence in SARS-CoV.

#### MHC class II results

We excluded peptides that have been observed with a mutation probability greater than 0.001 or are predicted to have non-zero probability of being glycosylated. We use the predicted binding affinity from NetMHCIIpan-4.0 for optimization and initial evaluation. With OptiVax-Unlinked, we design a vaccine with 19 peptides that achieves 91.67% EvalVax-Unlinked population coverage (averages over 15 geographic regions). As shown in Figure 5, the 19 vaccine peptides bind to a diverse range of alleles across the HLA-DRB/DP/DQ loci. Even though less effective than OptiVax-Robust on providing a high predicted number of average peptide-HLA hits in the population, the OptiVax-Unlinked peptide set still achieves high coverage on EvalVax-Robust metrics (93.23% for *p*(*n* ≥ 1), 70.19% for *p*(*n* ≥ 5), 45.87% for *p*(*n* ≥ 8)). The expected per-individual number of peptide-HLA hits for the design is 9.736, 8.454, and 6.860 for White, Black, and Asian populations, respectively (Table 1).

**Figure 5.**
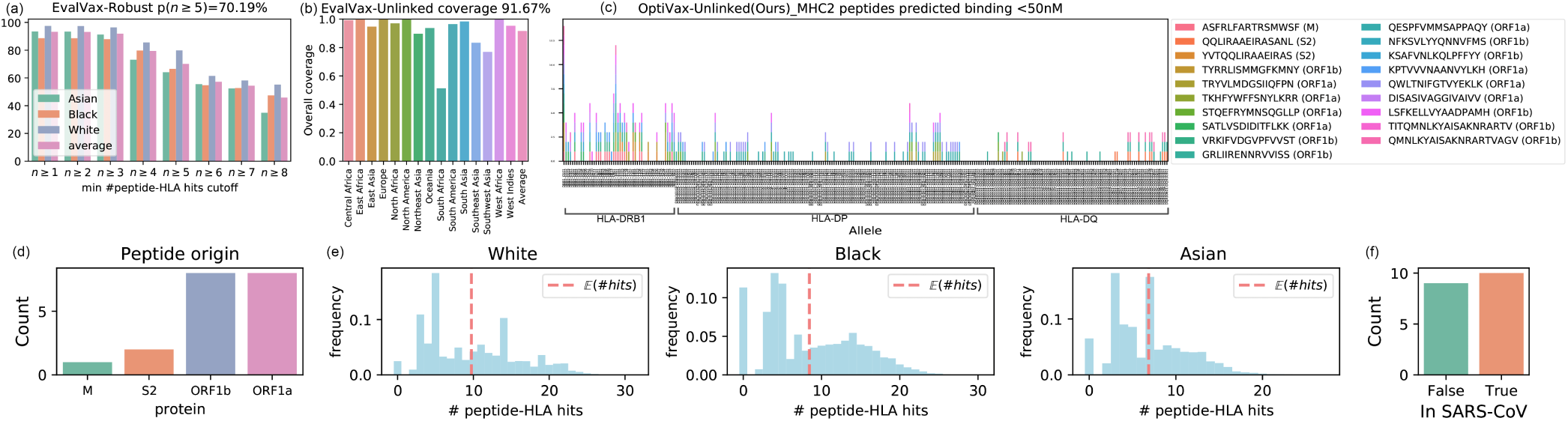
OptiVax-Unlinked selected optimal peptide set for MHC class II. (a) EvalVax-Robust population coverage at different minimum number of peptide-HLA hit cutoffs. (b) EvalVax-Unlinked population coverage. (c) Binding of vaccine peptides to 280 HLA-DRB1/DP/DQ alleles. (d) Distribution of peptide origin. (e) Distribution of the number of per-individual peptide-HLA hits in White/Black/Asian populations. (f) Peptide presence in SARS-CoV.

### 3.4 EvalVax evaluation of public vaccine designs for SARS-CoV-2

We used EvalVax to evaluate peptide vaccines proposed by other publications [52, 24, 53, 54, 55, 56, 57, 58, 59, 60, 61, 62, 63, 64, 65, 66, 67, 68, 69] on metrics including EvalVax-Unlinked and EvalVax-Robust population coverage at different per-individual number of peptide-HLA hits thresholds, expected per-individual number of peptide-HLA hits in White, Black, and Asian populations, percentage of peptides that are predicted to be glycosylated, peptides observed to mutate with greater than 0.001 probability, or peptides that sit on known cleavage sites. We define *vaccine efficiency* as the mean expected per-individual number of peptide-HLA hits for a vaccine divided by the number of peptides in the vaccine. This metric represents the mean probability of display of each peptide in a vaccine, and normalizes vaccine performance by vaccine peptide count.

Figures 6 to 9 show the comparison between OptiVax-Robust designed MHC class I and class II vaccines at all vaccine sizes (top solution in the beam up to the given vaccine size) from 1-35 peptides (blue curves) and baseline vaccines (red crosses) proposed by other publications. We observe superior performance of OptiVax-Robust designed vaccines on all evaluation metrics at all vaccine sizes for both MHC class I and class II. Most baselines achieve reasonable coverage at *n* ≥ 1 peptide hits. However, many fail to show a high probability of higher hit counts, indicating a lack of predicted redundancy if a single peptide is not displayed. We also evaluate randomly selected peptide sets of size 19 from predicted binders of MHC class I and II, where a binder is defined as a peptide that is predicted to bind with ≤ 50nM to more than 5 of the alleles in the MHC class. We found that a random binder set can achieve coverage that outperforms some of the proposed vaccines that we use as baselines.

**Figure 6.**
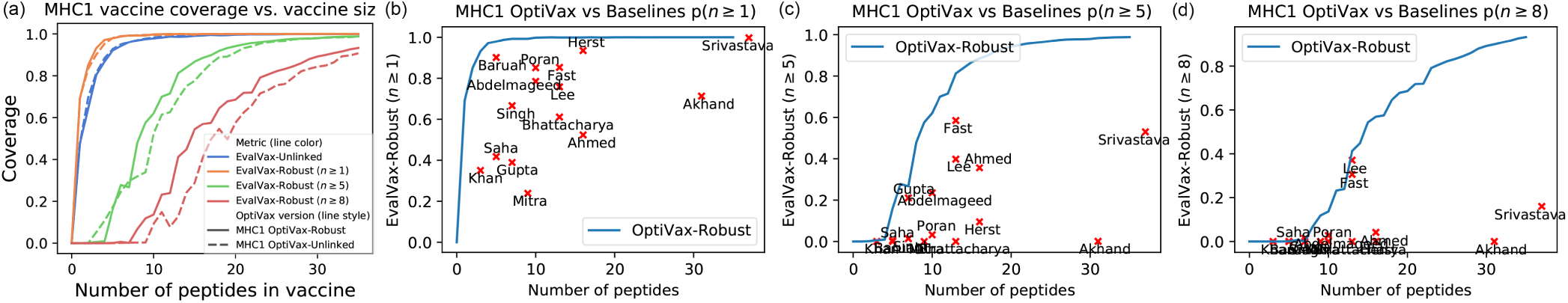
EvalVax population coverage evaluation for MHC class I vaccines. (a) EvalVax population coverage for OptiVax-Unlinked and OptiVax-Robust proposed vaccine at different vaccine size (b) EvalVax-Robust population coverage with *n* ≥ 1 peptide-HLA hits per individual, OptiVax-Robust performance is shown by the blue curve and baseline performance is shown by red crosses (labeled by first author’s name) (c) EvalVax-Robust population coverage with *n* ≥ 5 peptide-HLA hits. (d) EvalVax-Robust population coverage with *n* ≥ 8 peptide-HLA hits.

**Figure 7.**
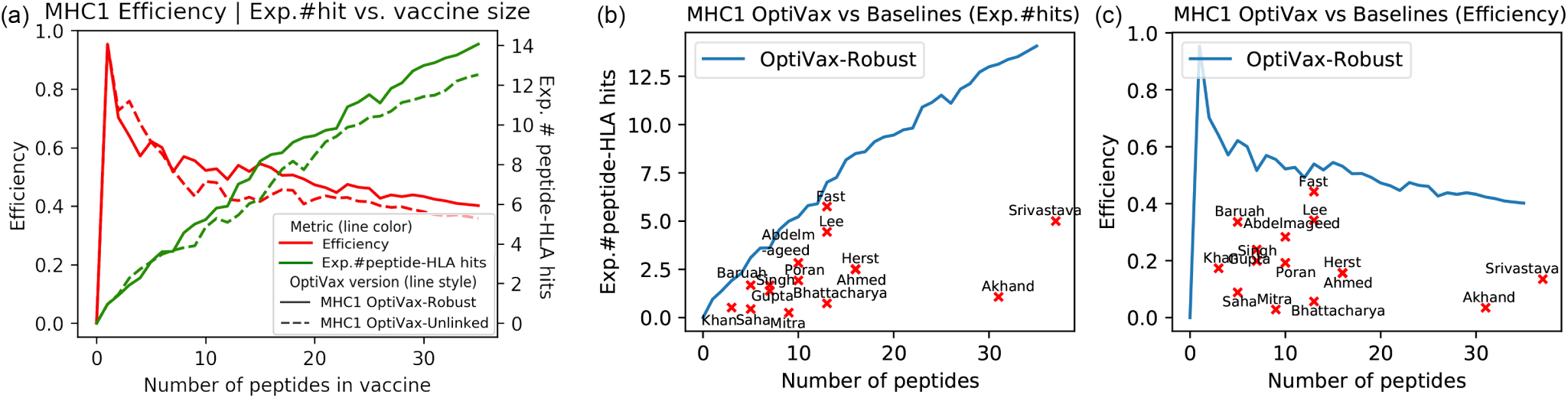
Expectation of per individual number of peptide-HLA hits and vaccine efficiency for MHC class I vaccines. (a) Expected number of peptide-HLA hits vs. peptide vaccine size for OptiVax-Robust and OptiVax-Unlinked, and efficiency (hits / vaccine size) at different vaccine size. (b) Comparison between OptiVax-Robust and baselines on expected number of peptide-HLA hits. OptiVax-Robust performance is shown by the blue curve and baseline performance is shown by red crosses (c) Comparison between OptiVax-Robust and baselines on efficiency.

**Figure 8.**
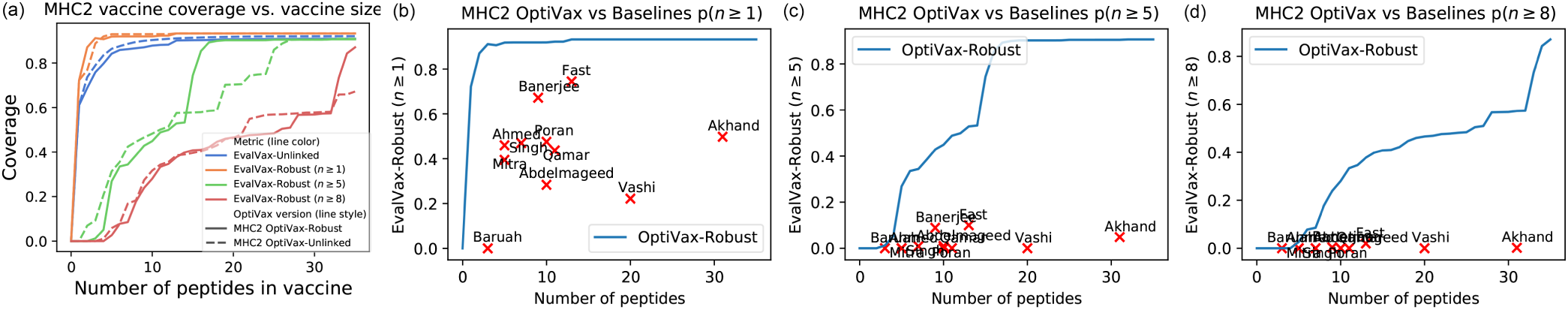
EvalVax population coverage evaluation for MHC class II vaccines. (a) EvalVax population coverage for OptiVax-Unlinked and OptiVax-Robust proposed vaccine at different vaccine sizes. (b) EvalVax-Robust population coverage with *n* ≥ 1 peptide-HLA hits per individual, OptiVax-Robust performance is shown by the blue curve and baseline performance is shown by red crosses (labeled by first author’s name). (c) EvalVax-Robust population coverage with *n* ≥ 5 peptide-HLA hits. (d) EvalVax-Robust population coverage with *n* ≥ 8 peptide-HLA hits.

**Figure 9.**
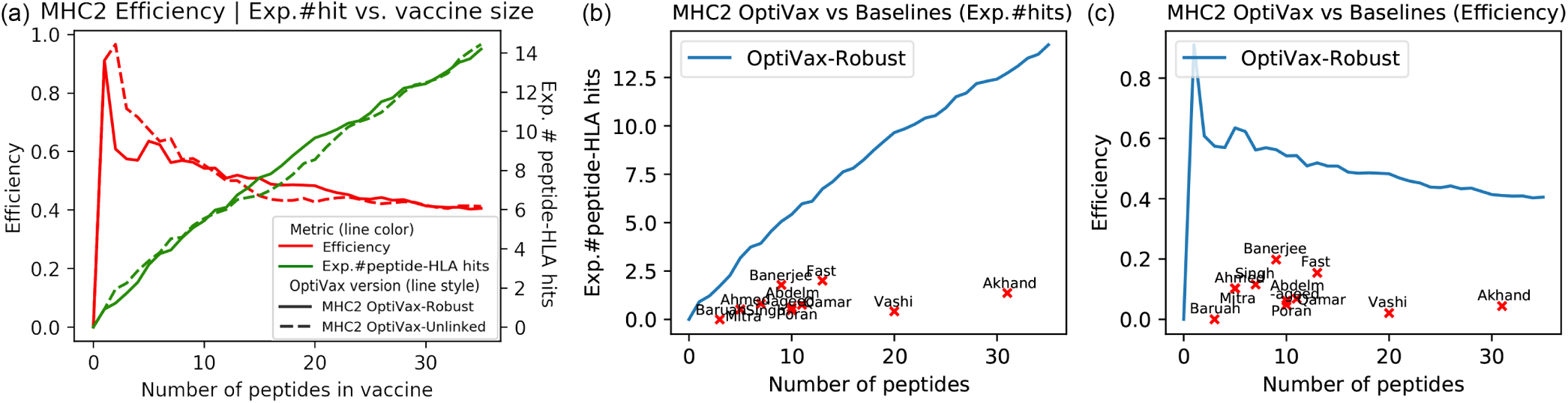
Expectation of per individual number of peptide-HLA hits and vaccine efficiency for MHC class II vaccines. (a) Expected number of peptide-HLA hits vs. peptide vaccine size for OptiVax-Robust and OptiVax-Unlinked, and efficiency (hits / vaccine size) at different vaccine size. (b) Comparison between OptiVax-Robust and baselines on expected number of peptide-HLA hits. OptiVax-Robust performance is shown by the blue curve and baseline performance is shown by red crosses. (c) Comparison between OptiVax-Robust and baselines on efficiency.

Table 1 summarizes EvalVax results for all baselines with a vaccine peptide count less than 150 peptides. We also included evaluation on peptide sets derived from taking all sliding windows with proper size for MHC class I and II from the S protein or S1 subunit, and evaluated an average of 500 random designs for MHC class I or class II that are comprised of 19 peptides that are predicted to bind either MHC class I and II. We found that the baseline methods all provide less coverage than OptiVax derived sets, and some contain peptides predicted to be glycosylated or have a high observed mutation probability (Table 1). We also observe some baselines contain peptides that sit on the cleavage sites or overlap with self-peptides. In addition, we found that for class II MHC coverage the S protein alone is unable to achieve more than 88% coverage for *n* ≥ 0 and 75.9% coverage *n* ≥ 5.

### 3.5 EvalVax results are robust to different binding prediction models

We evaluated all Table 1 vaccine designs using eleven independent peptide-MHC binding prediction methods to ensure that the performance observed in Table 1 is not an artifact. For MHC class I prediction we validated using seven methods: NetMHCpan-4.0; NetMHCpan-4.1; MHCflurry 1.6.0; PUFFIN; the mean of NetMHCpan-4.0 and MHCflurry 1.6.0 with a 50nM cutoff on predicted affinity; and NetMHCpan-4.0 and NetMHCpan-4.1 with a 99.5% cutoff on EL ranking. For MHC class II peptide-MHC binding prediction we validated using four different methods: NetMHCIIpan-3.2 and NetMHCIIpan-4.0, each with either a 50nM cutoff on predicted affinity or a 98% cutoff on EL ranking. The result of all eleven EvalVax evaluation metrics for all Table 1 designs are shown in Supplement Section S3. We find that all of the eleven methods we use for evaluation show that Table 1 is a conservative estimate of vaccine performance.

## 4 Discussion

The computational design of peptide vaccines for eliciting cellular immunity is built upon the imperfect science of predicting peptide presentation by MHC molecules. Peptide vaccine designs also need to ensure that individuals with rare MHC alleles display vaccine peptides to ensure a high rate of vaccine efficacy over the entire population.

To mitigate computational model uncertainty we have taken a very conservative view of peptide presentation, emphasizing precision over recall. To provide coverage for individuals with rare HLA types we use haplotype frequencies that include these types in our evaluations. We provide an evaluation tool, EvalVax, to permit the flexible analysis of vaccine proposals on key metrics, including population coverage and the expected number of peptides displayed. Not surprisingly, our OptiVax vaccine designs that are optimized with respect to EvalVax objective functions do well on the same metrics. We also find that OptiVax designs do well when evaluated on eleven computational models of peptide MHC binding, providing encouragement that their component peptides will be displayed.

EvalVax can be used for vaccine designs that are focused on the expression of viral proteins or their subunits to evaluate the level of viral peptide MHC presentation that is predicted to result. We note for SARS-CoV-2 in Table 1 that S protein and the S1 subunit both are limited in their predicted ability to provide robust population coverage for MHC class II display of more than five viral epitopes. This suggests that vaccines that only employ the S protein or its subunits may require additional peptide components for reliable CD4+ T cell activation across the entire population.

At present the World Health Organization lists 79 COVID-19 vaccine candidates in clinical or preclinical evaluation [70], and the precise designs of most of these vaccines are not public. We encourage the early publication of vaccine designs to enable collaboration and rapid progress towards safe and effective vaccines for COVID-19.

All of our software and data are freely available as open source to allow others to use and extend our methods.

## Supporting information

GISAID Acknowledgements

## Acknowledgements

Michael Birnbaum, Brooke Huisman, and Jonathan Krog provided helpful discussions. Viral sequences are from GISAID (see acknowledgement spreadsheet). This work was supported by in part by Schmidt Futures and NIH grant R01CA218094 to D.K.G.

This project has been funded in part with federal funds from the Frederick National Laboratory for Cancer Research, under Contract No. HHSN261200800001E. The content of this publication does not necessarily reflect the views or policies of the Department of Health and Human Services, nor does mention of trade names, commercial products, or organizations imply endorsement by the U.S. Government. This Research was supported in part by the Intramural Research Program of the NIH, Frederick National Lab, Center for Cancer Research. The views expressed in this article do not necessarily reflect the official policy or position of the National Institutes of Health, the Department of the Navy, the Department of Defense, or any other agency of the US government.

## Data and Software Availability

Our data and code are available at: https://github.com/gifford-lab/optivax

## Supplementary Information

### S1 Validation of Computational Peptide-MHC Prediction Models

#### S1.1 Criteria for Predicted Binding

NetMHCpan-4.0 [18] and NetMHCIIpan-4.0 [36] output predicted binding affinity (BA), percentile rank of predicted BA compared to a set of random natural peptides, and percentile rank of an eluted ligand (EL) score compared to a set of random natural peptides. Default parameters for these methods suggest EL percentile rank thresholds of 0.5% and 2% rank for classifying peptides as strong and weak binders, respectively, for MHC class I and thresholds of 2% and 10% for strong and weak binders, respectively, for MHC class II.

To identify binders for our vaccine designs, we use a 50nM predicted binding affinity threshold (Section 2.2). We find binders selected with this criterion are also considered binders under alternative criteria based on percentile rank. Across our set of all candidate SARS-CoV-2 MHC class I peptides (Section 2.1), we find that 91.0% of peptide-MHC hits with ≤ 50nM predicted binding affinity by NetMHCpan-4.0 are also considered binders using BA percentile rank ≤ 0.5% (100.0% have BA percentile rank ≤ 2%). Using percentile rank for EL scores, 67.6% of peptide-MHC hits with ≤ 50nM predicted binding affinity have EL percentile rank ≤ 0.5% (92.6% have EL percentile rank ≤ 2%). Across all candidate SARS-CoV-2 MHC class II peptides, we find that 86.1% of peptide-MHC hits with ≤ 50nM predicted binding affinity by NetMHCIIpan-4.0 are also considered binders using BA percentile rank ≤ 2% (100.0% have BA percentile rank ≤ 10%). Using percentile rank for EL scores, 26.1% of peptide-MHC hits with ≤ 50nM predicted binding affinity have EL percentile rank ≤ 2% (63.1% have EL percentile rank ≤ 10%).

#### S1.2 Validation on SARS-CoV-2 and SARS-CoV Experimental Data

We evaluate peptide-MHC binding predictions on a set of experimentally assessed SARS-CoV-2 peptides whose peptide-MHC complex stability was assessed in vitro across 11 MHC allotypes (5 HLA-A, 1 HLA-B, 4 HLA-C, 1 HLA-DRB1) [38]. Prachar et al. [38] suggest that peptides with low (< 60%) stability are unlikely to elicit an immune response and are unsuitable for vaccine development. For MHC class I alleles, the dataset contains 912 unique peptides-MHC pairs, of which 185 peptides are considered stable (≥ 60% stability). For MHC class II, the dataset contains 93 total peptides, of which 22 are stable. We use our computational models to predict peptide-MHC binding and evaluate them using various binding criteria against the experimental peptide stability measurement (Table S1). AUROC and average precision are computed using raw predictions, and the remaining metrics are computed using binarized predictions based on the respective binding criteria (using scikit-learn [71]). We compare classification performance using different binding criteria (see Section S1.1) and find in general that classifying binders using predicted binding affinity using a 50nM threshold maximizes AUROC and precision (Table S1). We find that our mean ensemble of NetMHCpan-4.0 and MHCflurry further improves classification AUROC and precision over the individual models for predicting MHC class I epitopes. On MHC class II data, we note NetMHCIIpan-4.0 achieves AUROC 0.848 and precision 0.625 using a 500nM threshold (Table S1). While NetMHCIIpan-4.0 with a 50nM threshold does not identify any peptides in this dataset as binders, we use this stricter threshold in our vaccine designs as it is more conservative and less likely to admit false positive binders. In general, we find performance of PUFFIN with a 50nM binding threshold comparable to alternative methods on both MHC class I and class II data and use PUFFIN as part of our vaccine design evaluation.

We additionally validate our computational models using previously reported SARS-CoV T cell epitopes from experimental studies [39, 40, 41] as provided by Fast et al. [24]. For MHC class I, this dataset contains 17 experimentally-determined HLA-A*02:01 associated CD8 T cell epitopes and 1236 non-epitope 9-mer peptides from the rest of the SARS-CoV Spike (S) protein. Table S2 shows the performance of our peptide-MHC binding prediction models on these SARS-CoV peptides.

**Table S1.** Classification performance of computational methods for predicting peptide-MHC binding evaluated on experimental SARS-CoV-2 peptide stability data across 11 MHC allotypes (5 HLA-A, 1 HLA-B, 4 HLA-C, 1 HLA-DRB1). Ensemble outputs the mean predicted binding affinity of NetMHCpan-4.0 and MHCflurry (see Section 2.2). (BA = binding affinity, EL = eluted ligand)

| MHC | Model | Binding Criterion | AUROC | Precision | Sensitivity | Specificity | Avg. Precision |
| --- | --- | --- | --- | --- | --- | --- | --- |
| Class I | NetMHCpan-4.0 | BA ≤ 50nM | 0.845 | 0.516 | 0.714 | 0.829 | 0.486 |
| | NetMHCpan-4.0 | BA ≤ 500nM | 0.845 | 0.308 | 0.968 | 0.446 | 0.486 |
| | NetMHCpan-4.0 | BA % Rank ≤ 0.5 | 0.746 | 0.249 | 0.968 | 0.257 | 0.416 |
| | NetMHCpan-4.0 | BA % Rank ≤ 2 | 0.746 | 0.212 | 1.000 | 0.054 | 0.416 |
| | NetMHCpan-4.0 | EL % Rank ≤ 0.5 | 0.757 | 0.256 | 0.930 | 0.312 | 0.479 |
| | NetMHCpan-4.0 | EL % Rank ≤ 2 | 0.757 | 0.214 | 0.989 | 0.077 | 0.479 |
| | NetMHCpan-4.1 | BA ≤ 50nM | 0.853 | 0.504 | 0.719 | 0.820 | 0.499 |
| | NetMHCpan-4.1 | BA ≤ 500nM | 0.853 | 0.304 | 0.984 | 0.428 | 0.499 |
| | NetMHCpan-4.1 | EL % Rank ≤ 0.5 | 0.776 | 0.278 | 0.903 | 0.403 | 0.490 |
| | NetMHCpan-4.1 | EL % Rank ≤ 2 | 0.776 | 0.219 | 0.989 | 0.103 | 0.490 |
| | MHCflurry 1.6.0 | BA ≤ 50nM | 0.724 | 0.404 | 0.422 | 0.842 | 0.411 |
| | PUFFIN | BA ≤ 50nM | 0.768 | 0.526 | 0.492 | 0.887 | 0.485 |
| | PUFFIN | BA ≤ 500nM | 0.768 | 0.272 | 0.870 | 0.406 | 0.485 |
| | Ensemble | Mean BA ≤ 50nM | 0.862 | 0.683 | 0.514 | 0.939 | 0.650 |
| Class II | NetMHCIIpan-4.0 | BA ≤ 50nM | 0.848 | 0.000 | 0.000 | 1.000 | 0.762 |
| | NetMHCIIpan-4.0 | BA ≤ 500nM | 0.848 | 0.625 | 0.682 | 0.873 | 0.762 |
| | NetMHCIIpan-4.0 | EL % Rank ≤ 2 | 0.908 | 1.000 | 0.182 | 1.000 | 0.785 |
| | NetMHCIIpan-4.0 | EL % Rank ≤ 10 | 0.908 | 0.789 | 0.682 | 0.944 | 0.785 |
| | NetMHCIIpan-3.2 | BA ≤ 50nM | 0.766 | 1.000 | 0.045 | 1.000 | 0.544 |
| | NetMHCIIpan-3.2 | BA ≤ 500nM | 0.766 | 0.253 | 0.909 | 0.169 | 0.544 |
| | NetMHCIIpan-3.2 | BA % Rank ≤ 2 | 0.766 | 0.380 | 0.864 | 0.563 | 0.536 |
| | NetMHCIIpan-3.2 | BA % Rank ≤ 10 | 0.766 | 0.253 | 1.000 | 0.085 | 0.536 |
| | PUFFIN | BA ≤ 50nM | 0.704 | 0.667 | 0.091 | 0.986 | 0.430 |
| | PUFFIN | BA ≤ 500nM | 0.704 | 0.275 | 0.864 | 0.296 | 0.430 |

**Table S2.**
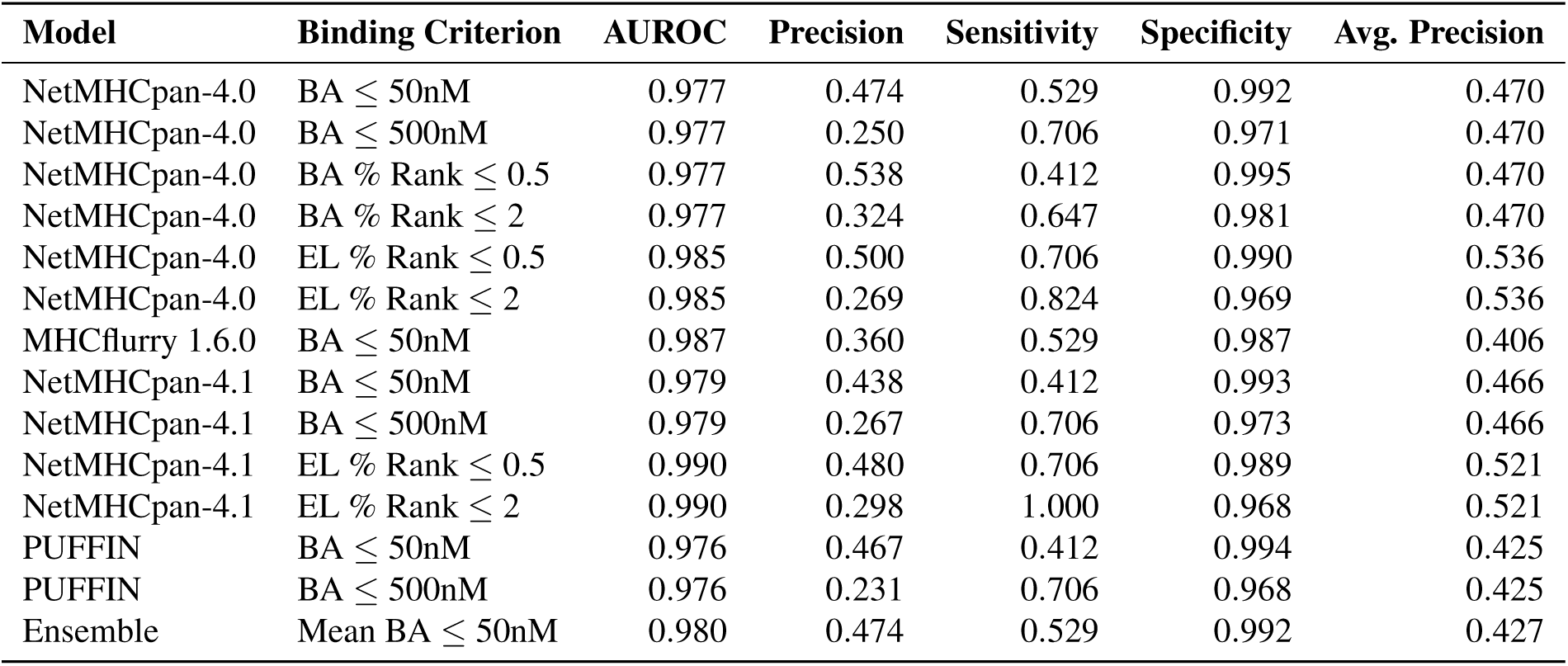
Classification performance of computational methods for predicting peptide-MHC binding evaluated on 17 experimentally determined HLA-A*02:01 associated CD8 T-cell epitopes from SARS-CoV vs. rest of SARS-CoV Spike (S) protein. Ensemble outputs the mean predicted binding affinity of NetMHCpan-4.0 and MHCflurry (see Section 2.2). (BA = binding affinity, EL = eluted ligand)

| Model | Binding Criterion | AUROC | Precision | Sensitivity | Specificity | Avg. Precision |
| --- | --- | --- | --- | --- | --- | --- |
| NetMHCpan-4.0 | BA ≤ 50nM | 0.977 | 0.474 | 0.529 | 0.992 | 0.470 |
| NetMHCpan-4.0 | BA ≤ 500nM | 0.977 | 0.250 | 0.706 | 0.971 | 0.470 |
| NetMHCpan-4.0 | BA % Rank ≤ 0.5 | 0.977 | 0.538 | 0.412 | 0.995 | 0.470 |
| NetMHCpan-4.0 | BA % Rank ≤ 2 | 0.977 | 0.324 | 0.647 | 0.981 | 0.470 |
| NetMHCpan-4.0 | EL % Rank ≤ 0.5 | 0.985 | 0.500 | 0.706 | 0.990 | 0.536 |
| NetMHCpan-4.0 | EL % Rank ≤ 2 | 0.985 | 0.269 | 0.824 | 0.969 | 0.536 |
| MHCflurry 1.6.0 | BA ≤ 50nM | 0.987 | 0.360 | 0.529 | 0.987 | 0.406 |
| NetMHCpan-4.1 | BA ≤ 50nM | 0.979 | 0.438 | 0.412 | 0.993 | 0.466 |
| NetMHCpan-4.1 | BA ≤ 500nM | 0.979 | 0.267 | 0.706 | 0.973 | 0.466 |
| NetMHCpan-4.1 | EL % Rank ≤ 0.5 | 0.990 | 0.480 | 0.706 | 0.989 | 0.521 |
| NetMHCpan-4.1 | EL % Rank ≤ 2 | 0.990 | 0.298 | 1.000 | 0.968 | 0.521 |
| PUFFIN | BA ≤ 50nM | 0.976 | 0.467 | 0.412 | 0.994 | 0.425 |
| PUFFIN | BA ≤ 500nM | 0.976 | 0.231 | 0.706 | 0.968 | 0.425 |
| Ensemble | Mean BA ≤ 50nM | 0.980 | 0.474 | 0.529 | 0.992 | 0.427 |

### S2 Details on S, M, N protein only vaccine design

**Figure S1.**
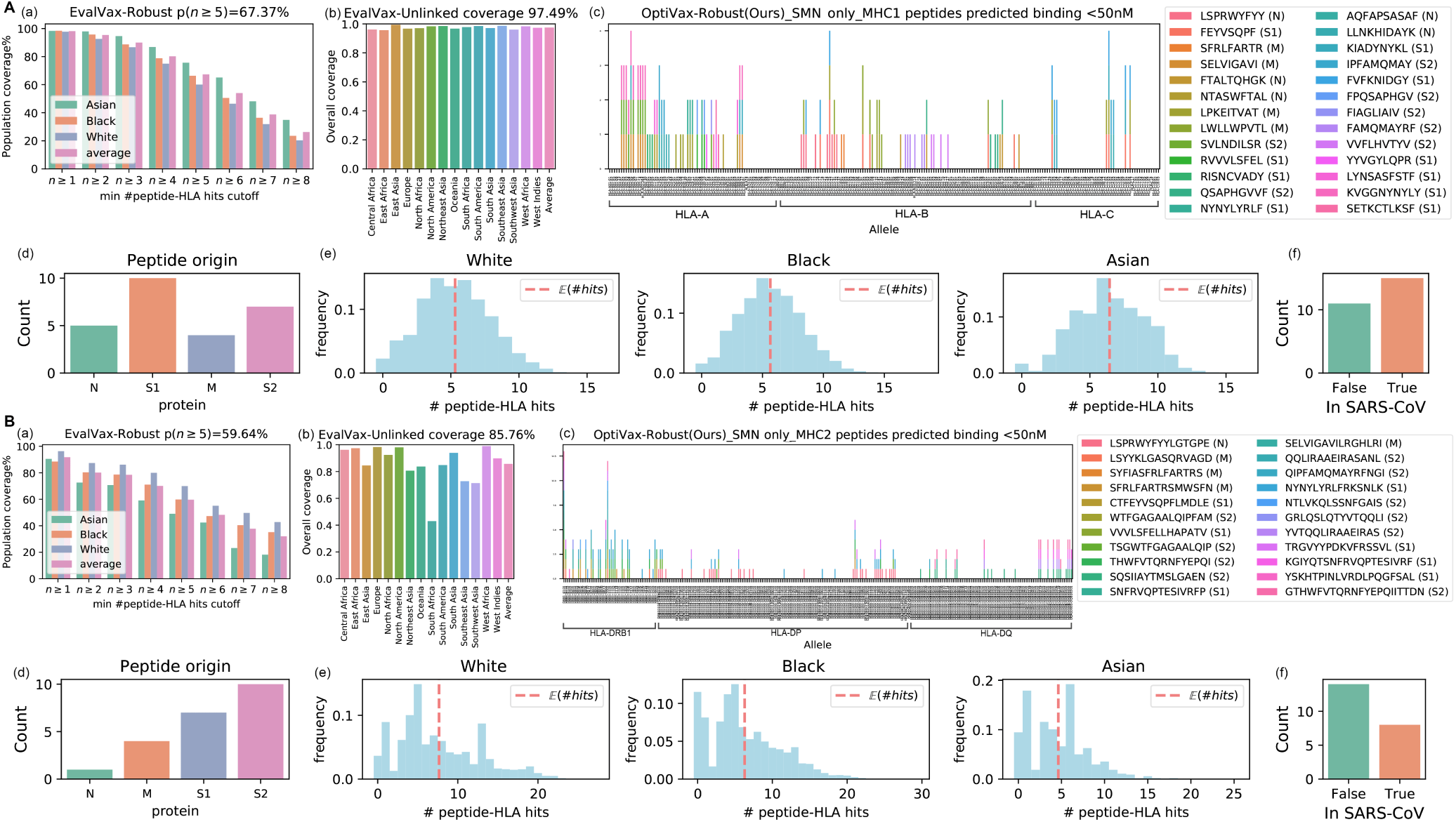
OptiVax-Robust designed vaccine using peptides from S, M, and N proteins only. (A) Results for MHC class I. (B) Results for MHC class II. (a) EvalVax-Robust population coverage at different minimum number of peptide-HLA hit cutoffs. (b) EvalVax-Unlinked population coverage. (c) Binding of vaccine peptides to each of the available alleles in MHC I and II. (d) Distribution of peptide origin. (e) Distribution of the number of per-individual peptide-HLA hits in White/Black/Asian populations. (f) Peptide presence in SARS-CoV.

### S3 Evaluation of baseline and OptiVax vaccines using different prediction tools/binder calling strategies

See supplementary table in `Supplementary_S3_evaluation_on_different_tools.xlsx`.

### S4 Detailed GISAID accessions

See table in `GISAID_Acknowledgements.xlsx` for acknowledgements.

